# Uncovering the dynamic mechanisms of the *Pseudomonas aeruginosa* quorum sensing and virulence networks using Boolean modelling

**DOI:** 10.1101/756551

**Authors:** Manuel Banzhaf, Osbaldo Resendis-Antonio, M. Lisandra Zepeda-Mendoza

## Abstract

**Introduction:** *Pseudomonas aeruginosa* is an opportunistic pathogen with an extraordinary metabolic adaptability and a large repertoire of virulence factors that allow it to cause acute and chronic infections. Treatment of *P. aeruginosa* infections often fail due to its antibiotic resistance mechanisms, thus novel strategies aim at targeting virulence factors instead of growth-related features. However, there is currently not a clear understanding of the dynamic nature inherent to the wiring of its virulence networks.

**Results:** In this study, we manually reconstructed the signalling and transcriptional regulatory networks of 12 acute (incl. pyocin and elastase) and 8 chronic virulence factors (incl. biofilm), and the 4 quorum sensing (QS) systems of *P. aeruginosa.* Using Boolean modelling (BM), we unveiled the important roles that stochasticity and node connectivity play in the networks’ inherent dynamicity and robustness. We showed that both the static interactions, as well as the time when the interactions take place, are important features in the QS network. In addition, we found that the virulence factors of the acute networks are under strict repression, or under an activation that is non-strict or oscillatory, while the chronic networks favour the repression of the virulence factor, with only moderate activation under certain conditions.

**Conclusion:** In conclusion, our *in silico*-modelling framework provided us with a system-level view of the *P. aeruginosa* virulence and QS networks to gain new insights into the various mechanisms that support its pathogenicity and response to stressors targeting these networks. Thus, we suggest that BM provides an invaluable tool to guide the design of new treatments against *P. aeruginosa.*

## Introduction

*Pseudomonas aeruginosa* is an ubiquitous Gram-negative bacterium, well known for its enormous adaptability and resistance to environmental stressors (including antimicrobials) [1]. With several pathogenic strains, *P. aeruginosa* can cause wound infections and catheter associate urinary tract infections [2] and is a major health concern to immunocompromised and cystic fibrosis patients [3]. Its ability to colonize a host is largely due to the production of various virulence factors. For example, production of elastase, which is a protease that degrades a major lung structural component called elastin, is related to the severity of acute infections, while the formation of adhesive communities on surfaces (i.e. biofilms) contributes to chronic infections [4]. One of the main mechanisms in which *P. aeruginosa* controls its gene expression to respond and adapt to the environment is quorum sensing (QS) [5]-[7]. The four QS systems in *P. aeruginosa* are Las, Rhl, *Pseudomonas* quinolone signal (PQS), and the integrated QS (IQS). They entail an intricate and highly interconnected regulatory network with a hierarchical architecture that controls the bacterial virulence as a function of its population density by detecting the amount of autoinducer (AI) molecules in the environment [5]. The QS network in *P. aeruginosa* is governed by the Las system, which produces and senses the AI N-3-oxo-dodecanoyl-L-Homoserine lactone [8] and activates the expression of multiple virulence genes such as those producing elastase [9]. The Las system also activates the Rhl system, which produces and senses the AI N-butyryl-L-Homoserine lactone [10]. In addition, the quinolone dependent system known as PQS is positively regulated by Las and negatively by Rhl [11], and it activates the Rhl system to regulate itself. Recently, the IQS system was found to sense 2-(2-hydroxyphenyl)-thiazole-4-carbaldehyde to also integrate stress cues with the QS network [12].

The rise of antibiotic resistance in *P. aeruginosa* and the slow entry of new drugs to market pose a serious risk to human health. As an alternative drug development strategy, instead of targeting growth-related features (e.g. membrane, DNA and protein synthesis), it has been proposed to target the virulence factors and the QS network [13]. Compared to a traditional antimicrobial therapy, this approach might be advantageous, as there is no selection pressure on the bacteria and the infectious capacity is diminished, allowing the host’s immune system to clear the infection. Different computational approaches, particularly genome-scale metabolic modelling [14], [15] and topological analyses [16], have been proven useful for identifying key elements that can be targeted to disrupt the network’s connectivity [17]. However, these approaches offer little insights into the dynamic responses of the regulatory and signalling networks, so that other modelling techniques need to be explored, such as Boolean modelling (BM).

In a Boolean model, the nodes represent the proteins, genes, or signalling metabolites that are part of the network, and the edges represent the activation or repression interactions between them. Each node can be in an ON (1, meaning active, expressed, or present) or OFF (0, meaning inactive, not expressed, or absent) state. At the beginning of the simulation, only the nodes defined as initial conditions are ON. The state of each node at a given time step (*t*(n)) in a simulation depends on their updating rule, which is defined by the basic logical operators AND, OR, and NOT. The more advanced operators THR and MOD can also be used. THR defines a lagging time for a node to have been active before executing its effect on another node, and MOD determines for how many time steps (*t*) the effect of one node upon another is executed. There are two modes in which the state of the nodes can be updated in each time step, called synchronous and asynchronous. In the synchronous mode, all the nodes are updated at the same time and the state of each node at *t*(n) depends on the states of their regulator nodes at *t*(n-1). This means that the network will always reach the same state for a given *t*(n) and a set of initial conditions. In the asynchronous mode, the updating order of the nodes is selected randomly during each iteration and they update according to the last update of their regulator nodes [18]. This mode induces complexity and variability into the system, so that the same initial conditions can result in different final states for a given *t*(n).

BM can study the activation dynamics between elements in interconnected biological networks [19], [20] and has been successfully applied to understand pathogenesis. However, it has mainly been used on the human immune system [21], with few studies focused on understanding the bacterial pathogenic networks. For example, a BM study on the Las, Rhl, and PQS systems found that the regulatory wiring of the QS network plays a dominant role in its robustness, suggesting that targeting the elements involved in QS is a promising strategy for drug development [22]. Although several efforts have been placed in targeting the QS [23], [24], the effect of the abundance of the AI in the environment and the effect of the lag time between the regulation of the nodes, have not been deeply examined with computational approaches. Most importantly, QS can elicit network connectivity rewiring to readjust the bacterial metabolism [25] and it has been noted that it has the capacity for compensatory mutations [26], [27]. All of these aspects should be considered for targeted drug design.

Although BM provides a mathematical description of the dynamic capacities of the virulence signalling and regulatory networks of bacterial pathogens, studies implementing such methodologies are scarce. Prompted by the necessity of understanding the adaptability inherent to the wiring of the *P. aeruginosa* virulence networks, we developed Boolean models of the 4 QS systems, and of the main transcriptional regulatory and signalling interactions involved in 12 acute and 8 chronic virulence factors. Using BM techniques on these models, we characterized the effect of stochasticity and network perturbations upon the activation of the virulence factor. Also, we could classify the networks according to their mechanisms for controlling the activation of their virulence factor as *i*) strictly or non-strictly repressing, and *ii*) not strictly or oscillating activating. Taken together, our results provide a novel picture of the complexity and dynamicity of the pathogenicity networks of *P. aeruginosa*.

## Methods

### Construction of Boolean models

We performed an extensive literature review to define the nodes and updating rules of the networks representing acute and chronic virulence factors and the QS systems (Fig 1A). The QS network encompassed the Las, Rhl, PQS, and IQS systems [5], [28]. The acute virulence factors we studied were: the high-affinity iron-chelating compounds pyoverdine and pyochelin, pyocyanin (a compound that generates reactive toxic oxygen species), the antibiotic protein pyocin, exotoxin A (which inhibits protein synthesis), elastase, the secretion systems T1SS, T2SS, and T3SS, the motility related factors flagella and type IV pilli (TFP), and the Lysyl endopeptidase PrpL. The chronic virulence factors we studied were: the exopolysaccharide alginate (which protects the bacteria from environmental insults and enhances adhesion to solid surfaces), resistance to the antibiotics β-lactam and fluoroquinolone, the secretion systems T6SS-HIS-I/II/III, and biofilm formation (Suppl File 1).

**Figure 1.**
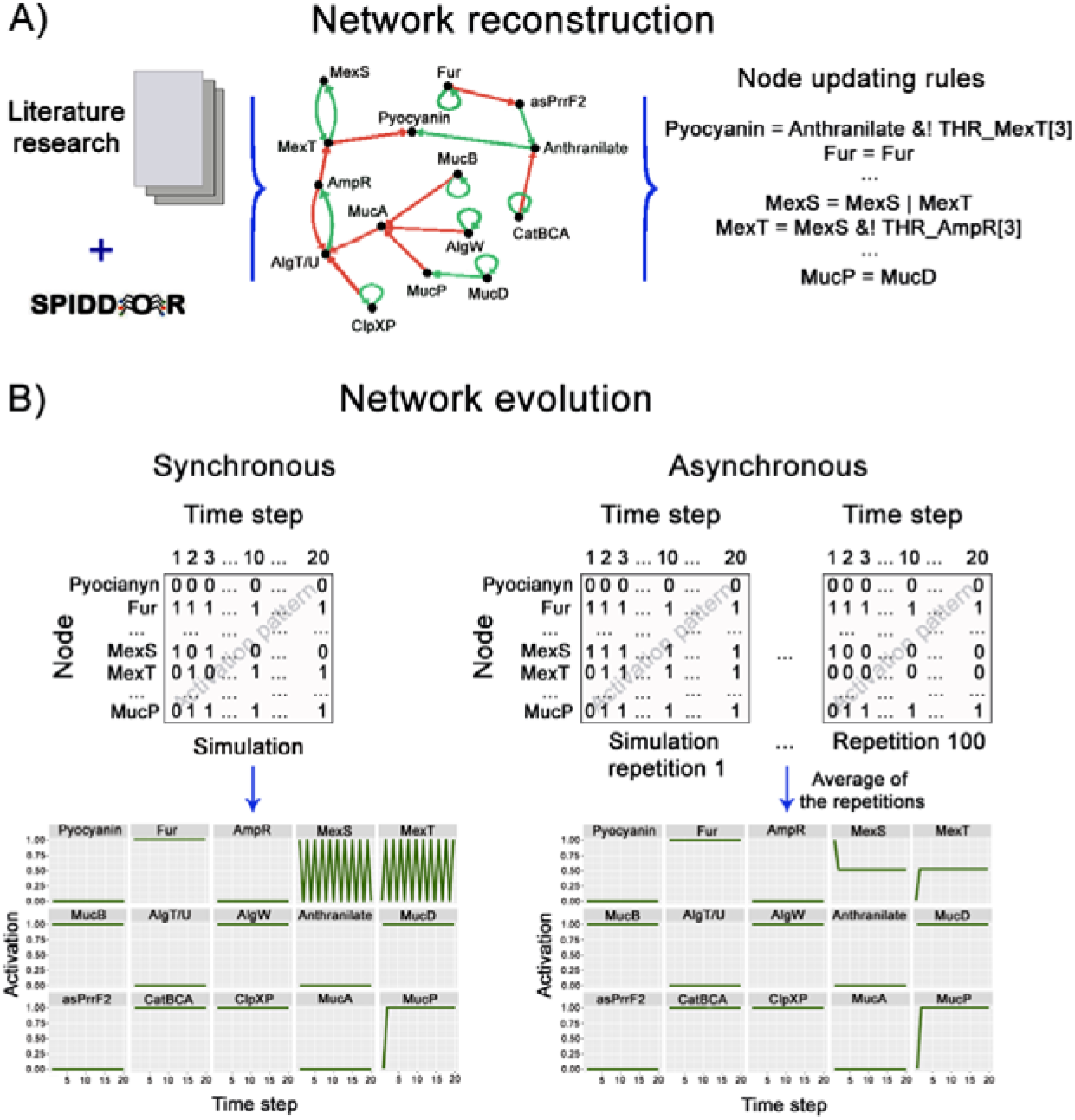
Schematic representation of the network reconstruction and analysis. A) We performed extensive literature research to identify the most relevant nodes (genes, metabolites, proteins) involved in the production of a virulence factor (e.g. pyocyanin). The nodes and their interactions were implemented into a Boolean model with the R package SPIDDOR, which allows for analyses of these interactions represented with Boolean updating rules. Nodes that are part of the set of initial conditions are defined as ON based on their own presence, for example, the Fur node, which is defined as Fur = Fur. B) The evolution of the state of each node overtime (i.e. the activation pattern of the network) can be analysed under the synchronous or the asynchronous updating modes. Since the synchronous mode is deterministic, only one simulation for each set of tested initial conditions is needed to find the activation patterns of the network nodes and a node can only have 1 or 0 activation values. On the other hand, several repetitions are needed for the stochastic asynchronous mode. The average of the activation values of the nodes across all the repetitions is obtained in order to define their activation patterns. In the plots showing the nodes activation patterns, the x-axis marks the *t*(n) along the simulation. For example, 10 marks the state of the node in middle of our 20-time steps simulation, and 20 marks the final state of the node in the simulation. The y-axis is the level of activation, for example, a value of 1 means that the node is ON in all the repetitions of the simulation, and 0.5 means that the node was ON in 50% of the repetitions of the simulation.

The R package SPIDDOR [20] was used for the network implementation and Boolean analyses, together with the package snowfall [29] for the parallelization of the computations. The networks were exported to the SBML format and deposited on the Cell Collective Repository [30] (see Suppl File 2 for the list of IDs).

### Boolean analyses

For all networks, we first used the dynamic_evolution function (20 time steps, 100 repetitions) to track the evolution of the state of each node over time to identify the activation pattern of the network, with particular attention on the nodes representing the virulence factor (Fig 1B). Then, we also identified the set of stable states (i.e. the attractors) that each network reaches from a given set of initial conditions, using the get_attractor function (20 time steps, 100 repetitions). To identify which nodes are key for the network functionality, we studied the network activation patterns after node perturbations (node deletions and lowering of node activity) with the function KO_matrix. This function defines a perturbation index of a node as the ratio of the activation of the node in the network after one of the nodes is eliminated from the network to the activation of the node when all the nodes are present. Thus, the result is a square matrix in which the number of rows and columns equals the number of nodes in the network and the values are the perturbation indexes. We made heat map plots for visualizing the perturbation results with the create_heatmap function, which transforms the matrix resulting from the perturbation analyses into ranked values prior to plotting.

As the synchronous mode sheds light on the complex behaviours that result purely due to the connectivity of the nodes in the system, while the asynchronous mode sheds light on the complexity that can arise due to the network dynamicity, we performed our analyses under both updating modes. For the asynchronous mode, we recorded the average times that each node was ON over all the simulation repetitions. To examine the effect of time lags on the regulation of specific nodes, we set different THR values to the specific nodes of interest. For example, a THR = 3 *t* means that the node must have been ON in the previous three iterations in order to affect its regulated node.

One of the QS-targeting approaches involves the use of molecules designed to outcompete the AIs for receptors [31], [32]. Thus, we also evaluated the number of time steps needed for the AI nodes of each QS system for the system to reach sustained activation. To this end, different MOD values were tested. For example, the initial condition representing the PQS AI with a MOD = 1 *t* means that the PQS AI node will only exert the effect on its regulated node for one time step. This represents the case in which the available PQS (i.e. PQS AI node being ON) is very little that it only lasts one iteration. This simulation reflects the condition in which the AI is available in the environment at minimal concentrations for the bacteria to take them, deplete them from the medium, and reach quorum (i.e. turn the system ON and synthesize its own AI).

All code is available through GitHub under the project name MLZM-lab/PA-BM.

## Results

### QS network

As the Las system is at the top of the hierarchical structure of the QS network (Suppl File 1), we first analysed how much Las AI is needed for the Las system to activate and influence the rest of the QS network. To this end, we tested different MOD values on the Las AI initial condition node and identified the activation pattern of the QS network. We saw that under the synchronous mode the Las system showed sustained activation with Las AI MOD >5 *t*, represented by the node Las_REG_ being ON (Suppl Fig 1). Under the asynchronous mode, it only needed Las AI MOD > 2 *t* to reach ~40% sustained activation (i.e. sustained activation in ~40% of the simulations), and Las MOD = 4 *t* to get 77% of sustained activation (Suppl Fig 1). Next, we evaluated the interplay between the activation of the rsaL node, which represses the Las system. As expected, when only the initial condition that activates rsaL was given, nothing except the rsaL module was ON. To investigate how the signals compete to regulate the Las system, we set both the Las AI with different MOD values and the rsaL activator as initial conditions. Under the synchronous mode with Las AI MOD >7 *t*, Las_REG_ was ON for a limited time, and PQS_REG_ for only a maximum of 2 *t*, with Rhl_REG_ reaching a sustained activation (Fig 2A). Notably, the asynchronous mode showed 80% of sustained activation for Rhl_REG_ with Las Al MOD = 6 *t*. When Las AI MOD >7 *t*, Las_REG_ stayed active for long enough to transiently activate PQS_REG_, before it was inactivated by the Rhl system (Fig 2A). This means that, even with RsaL being ON, if there is enough AI in the environment, *P. aeruginosa* can exert its effect by activating the Rhl system.

**Figure 2.**
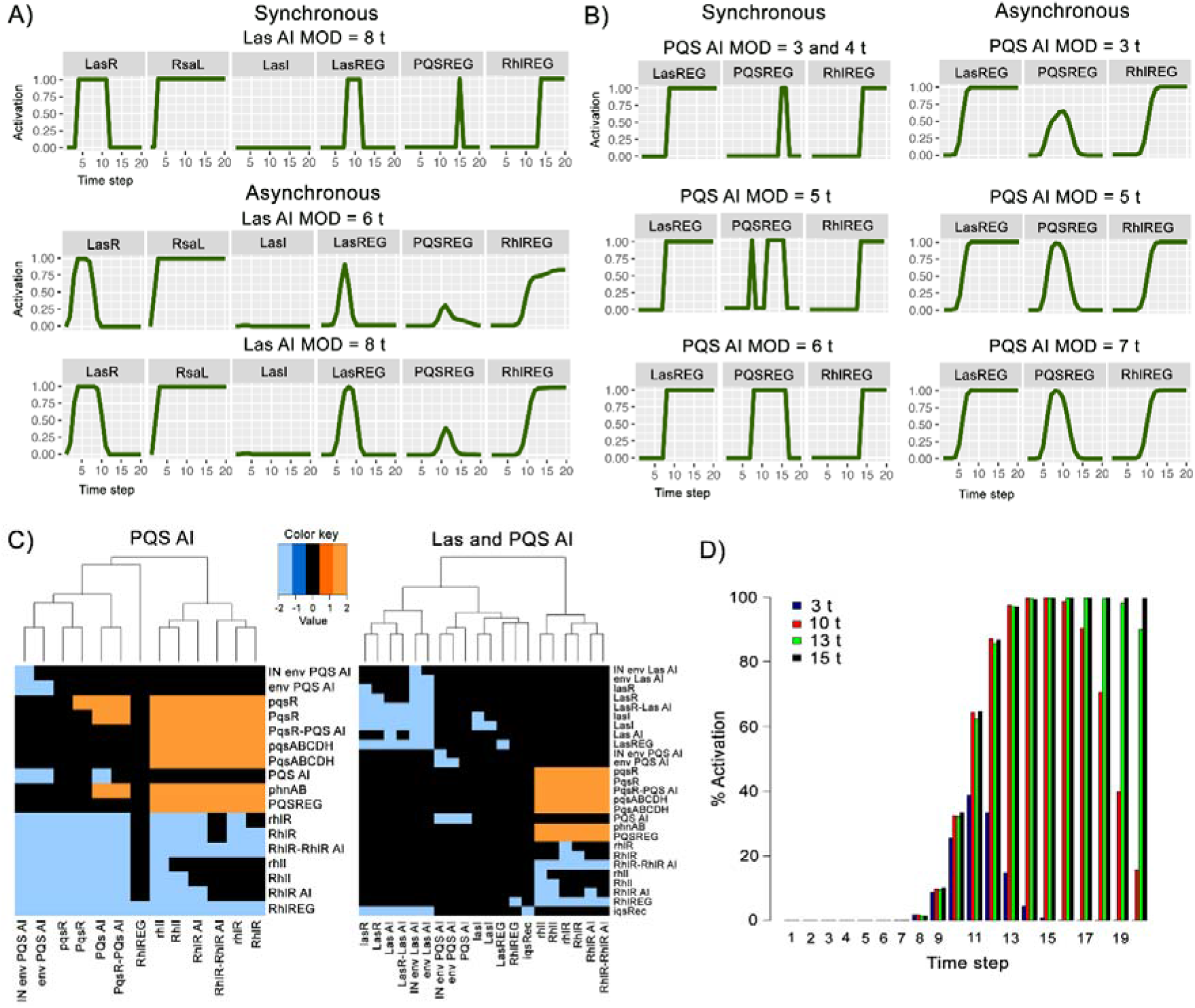
Activation patterns of the QS systems. A) Activation patterns of the Las system with the competing signals of Las AI and rsaL as initial conditions under the synchronous (first top row) and asynchronous updating modes with different Las AI MOD values (two bottom rows). B) Activation patterns of the PQS system with Las AI MOD = 6 *t* and PQS AI at various MOD values under the synchronous (left side) and asynchronous (right side) updating modes. C) Heat map of the rank values of the effect of node deletions on the QS network when in the environment there are PQS AI (left side) and PQS together with Las AIs (right side) under the asynchronous mode. The colour indicates if the node knockout entails a lower (blue) or higher (orange) activation of a component compared to an unperturbed simulation. D) Effect on the PQS system with the asynchronous mode when the repression by the Rhl system is delayed (THR = 3-15 *t*) and there is enough Las Al fora sustained activation of the Las system.

Since the PQS system is regulated by the Las, Rhl and IQS systems, we then aimed to identify which of these interactions are key and which are the conditions that influence them the most. To this end, we first tracked the evolution of the state of each node of the QS network over time under different initial conditions that directly influence the PQS system. We first set as initial conditions Las AI MOD = 6 *t* and PQS AI at various amounts (Fig 2B). In the synchronous mode, we saw that PQS AI with a short MOD of 3 and 4 *t* transiently activated the PQS system at *t*(15, 16). When the PQS AI MOD = 5 *t,* the PQS_REG_ showed two activation peaks, at *t*(8) and *t*(12-16). PQS_REG_ reached a constant activation when AI MOD > 5 *t*, however this activation was never sustained until the end of the simulation (*t*(20)) due to the repression by the Rhl system. This suggests that the most effective way of extending the activation of the PQS system is by modulating the activity of the Rhl system, not by modulating the PQS AI abundance. On the other hand, the asynchronous simulations showed PQS_REG_ with as much as ~50% of activation at *t*(8), and a maximum of ~67% at *t*(10) with PQS AI MOD = 3 *t*. As the PQS AI availability increased, PQS_REG_ was active in a higher percentage of the repetitions, but restricted to the *t*(6-12) in a normal shaped distribution of percentages of activation (with a 1^st^ and 3^rd^ quartiles of ~35%, and a maximum of ~99.8%). Thus, the asynchronous simulations represent the restriction in which the PQS system operates, but also the relative ease in which it can be activated in at least ~50% of the simulations, which can be considered to represent an activation in ~50% of the members in a population.

We then examined what would be the PQS system activation pattern if in the environment there were only various levels of PQS AI (Suppl Fig 2). To this end, we tested different PQS AI MOD values and evaluated the number of time steps needed for the PQS_REG_ node to reach sustained activation. With the synchronous mode, we found that a PQS AI MOD > 4 *t* was needed to activate PQS_REG_ for only one *t,* and a PQS AI MOD > 5 *t* for a PQS_REG_ sustained activation until it was turned OFF by the Rhl system that the PQS system activated for self-regulation. Notably, the asynchronous mode showed that the PQS system can get activated with a PQS AI MOD as short as 3 *t,* although the maximum activation was ~40.5% at *t*(9). These results confirm that the QS is wired in such a way that PQS can be easily activated but remain under strict regulation. Next, we evaluated how the node deletions and various amounts of AI for the different QS systems can affect the PQS system. We saw that the system was less affected by node deletions when there was only PQS AI than when there were also Las AIs present (Fig 2C). When more QS systems were included as active in the model, more nodes were affected by the perturbations. This was evident in the perturbation analyses with available AIs of the Las, Rhl, PQS, and IQS systems (Suppl Fig 3). Interestingly, although more nodes were affected when more systems were active, the QS network kept most of its nodes unaffected, with the changes restricted to the specific systems with the deleted node.

Lastly, we examined deeper the observation that PQS_REG_ was ON very briefly when the negative regulation by the Rhl system upon the PQS system had a THR = 3 *t*. We hypothesized that it is not only the hard-wired connections what shape the activation patterns of the nodes, but also the time in which these activations take place. To test this, we defined different THR values for the Rhl system regulation upon the PQS system. We delayed the repression by setting higher THR values and defined the Las AI at enough level for the Las system to reach a sustained activation (Las AI MOD = 6 *t*). As expected, in the synchronous mode, delaying the repression of the Rhl system on the PQS system for one *t* allowed the activation of PQS_REG_ one *t* longer, reaching a sustained activation when THR >6 *t* (Suppl Table 1). Notably, in the asynchronous mode with THR >7 *t,* PQS_REG_ reached activation of > 90% for more than one *t*, however it never reached 100% activation (Suppl Table 1, Fig 2D). Furthermore, the >90% activation was sustained only when THR > 12 *t.* To further explore the negative effect of the Rhl system on the PQS system, we evaluated the effect of the delay in the activation of the Rhl system by the Las system. Under the synchronous mode, delaying the activation by one *t* kept PQS_REG_ ON for one more *t,* as expected. However, it did not reach a sustained activation, as the Rhl system was activated by the IQS system, so that PQS_REG_ was OFF in *t*(19, 20). This result illustrates the robustness of the QS network, where the IQS system complements the Las system. Notably, the asynchronous mode showed a tighter regulation, with the PQS_REG_ having < 6% of activation since *t*(15-20), regardless of how much the activation of the Rhl system was delayed. These observations further suggest that both the dynamic nature of the network and the wiring of its nodes are key elements in the QS behaviour.

### Acute virulence networks

Similar to the analysis workflow used for the QS model, we examined our 12 acute virulence networks to identify their main mechanistic behaviours and adaptation strategies upon perturbations. In the following section, we describe only a selection of networks that clearly showed the main identified characteristics. See Suppl File 3 for additional information on the other identified relevant networks not discussed in this main text.

- *Strict repression of the virulence factor*

We first characterized the activation pattern of the nodes of every network in order to track the evolution of the state of each node over time, particularly the nodes representing the virulence factor. We found that the network for flagella formation is a robust system prone to the repression of the virulence factor. Under the scenario where all the initial conditions in the network were active (Suppl File 1), the flagella was never active under either updating mode. Only the deletion of the BifA node, which is an initial condition, led to an increase in the activation of flagella (Suppl Fig 4A). When we examined pairwise combinations of node deletions, only the BifA-WspR and BifA-WspF led to flagella activation as the attractor state. Thus, although this network represents a tightly controlled repressive system, it can skilfully alter its attractor state for the production of the virulence factor by changing the initial condition of only one of its nodes.

- *Non-strict activation of the virulence factor*

We also evaluated the attractors of the network to find the set of states to which the network tends to evolve. We found that the attractors of the small four-node pyocin model (Suppl File 1) in both updating modes showed an early sustained activation for pyocin production when RecA was part of the active initial conditions. However, pyocin was ON only transiently when RecA was deleted or had reduced activity (Suppl Fig 4B), and the 50% reduction of activity or deletion of PrtN led to pyocin being OFF during all the simulation. Notably, activation of the pyocin node upon RecA deletion was recovered by deleting or reducing the activity of PrtR (Suppl Tables 2, 3, Fig 3A). Thus, although the attractor is for pyocin to be ON, the network can straightforwardly modulate the virulence factor by modifying the activation of the other nodes.

**Figure 3.**
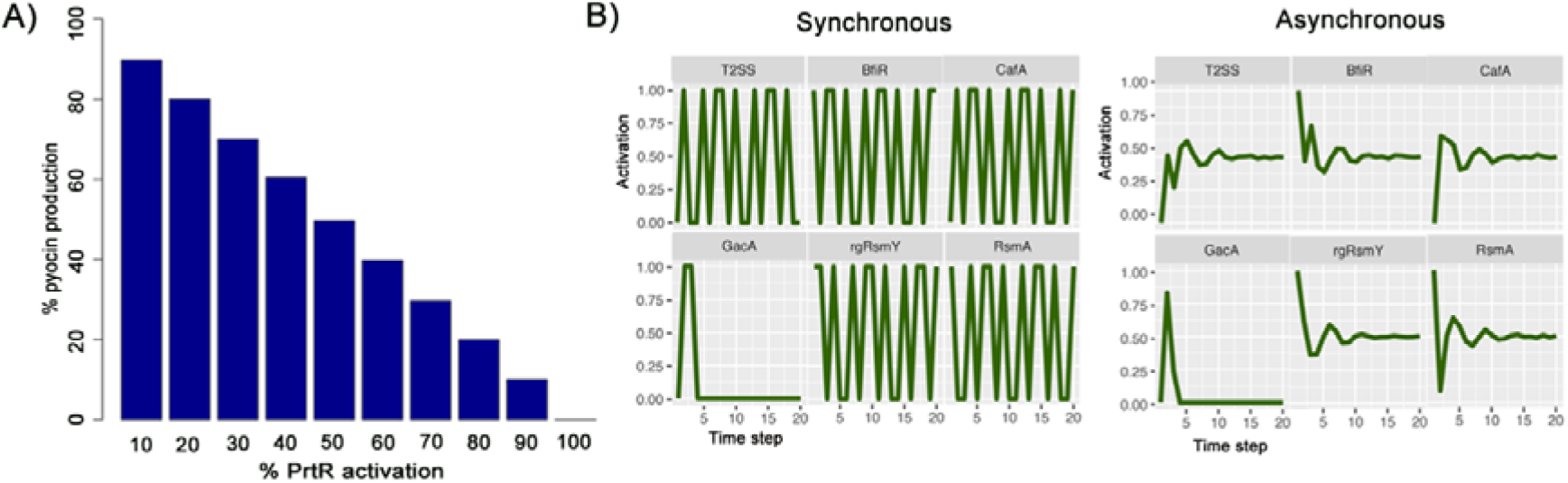
Activation patterns of the pyocin and T2SS networks. A) Activation of the pyocin node in a network with different levels of activity (ranging from 10 to 100%) of the node PrtR and all the initial conditions ON except RecA. B) Activation pattern of the T2SS network when the nodes CbrA and Vfr of the initial conditions directly activating the T2SS node were deleted under the synchronous (left side) and asynchronous (right side) updating modes.

- *Oscillatory activation of the virulence factor*

In our examination of the activation patterns of the nodes representing the virulence factors to track the evolution of their state over time, we identified an oscillatory activation pattern in some of the networks. For example, in the elastase production network (Suppl File 1) under the synchronous mode (Suppl Fig 5A). Notably, the asynchronous did not show an oscillating activation but a sustained 50% of activation of the elastase node (Suppl Fig 5B). This behaviour was determined by the interplay between the nodes MexS (an initial condition) and MexT (a repressor of elastase production), which activate each other. In the perturbation analyses, no single or pairwise node deletion led to less elastase (Suppl Fig 5C). Most notably, if any of the activities of the nodes was reduced to 50%, the elastase node was fully ON. We observed the same behaviour regardless of the combination of given initial conditions.

The T2SS network (Suppl File 1) also showed an oscillatory activation pattern when the nodes CbrA and Vfr of the initial conditions directly activating the T2SS were deleted (Fig 3B). Notably, although both updating modes had half of their attractors for activation and half for repression, the synchronous had 8 attractors with equal probability, while the asynchronous had 16 attractors, not all of them with the same probability (Suppl Table 4). These results suggest that the randomness in the network allows it to explore a wider space of responses and to rapidly adapt to an active state as soon as needed, in spite of the factor being metabolically expensive, such as the T2SS. Furthermore, when we deleted pairwise combinations of the initial condition nodes, the model responded with a third of the attractors leading to full inactivation, a third to full activation, and a third to 50% activation (Suppl Table 5). This highlights the network’s readiness to adapt to whatever behaviour the environment would require.

- *Stochasticity effect on the network attractor space*

We observed that a network does not need to be large (i.e. have many nodes) to exhibit interesting behaviours. Although small, the five-node model of pyoverdine production is complex enough to have negative regulations with delayed effects between the nodes (Suppl File 1). The synchronous mode showed two attractors, one with and one without pyoverdine production. Interestingly, the asynchronous mode also had these two attractors, but with unequal probabilities. The attractor with pyoverdine ON occurred in 99.8% of the simulations, while the one with it OFF happened in only 0.2% of the simulations. Thus, the asynchronous mode provided better insights into the network’s behaviour, which is more prone to pyoverdine production than to its repression.

Another interesting network from an energetically expensive factor is that of T3SS (Suppl File 1). The attractors of the system had T3SS OFF. However, the asynchronous mode showed a particular behaviour. Out of the 15 attractors, 7 led to an ON T3SS in only 1-3 (0.01%) of the simulations. To further explore this, we performed less repetitions (10-100 repetitions). Each analysis showed the same result, where 8 of the 16 attractors found in total had T3SS ON and occurred with an average probability of 0.008% (Suppl Table 6). This suggests that although T3SS is a tightly repressed factor, the attractor space of the network allows for the possibility of the production of the factor simply by stochasticity. Importantly, the results of the perturbation analyses under both updating modes are not the same (Fig 4). In the synchronous mode, the deletion of only two nodes (PtrB and RecA) positively affected the T3SS and none affected it negatively, while in the asynchronous mode, three node deletions (PtrB, RecA, and Crc) affected it positively and 31 affected it negatively. This further supports the importance of the effect of stochasticity upon the network.

**Figure 4.**
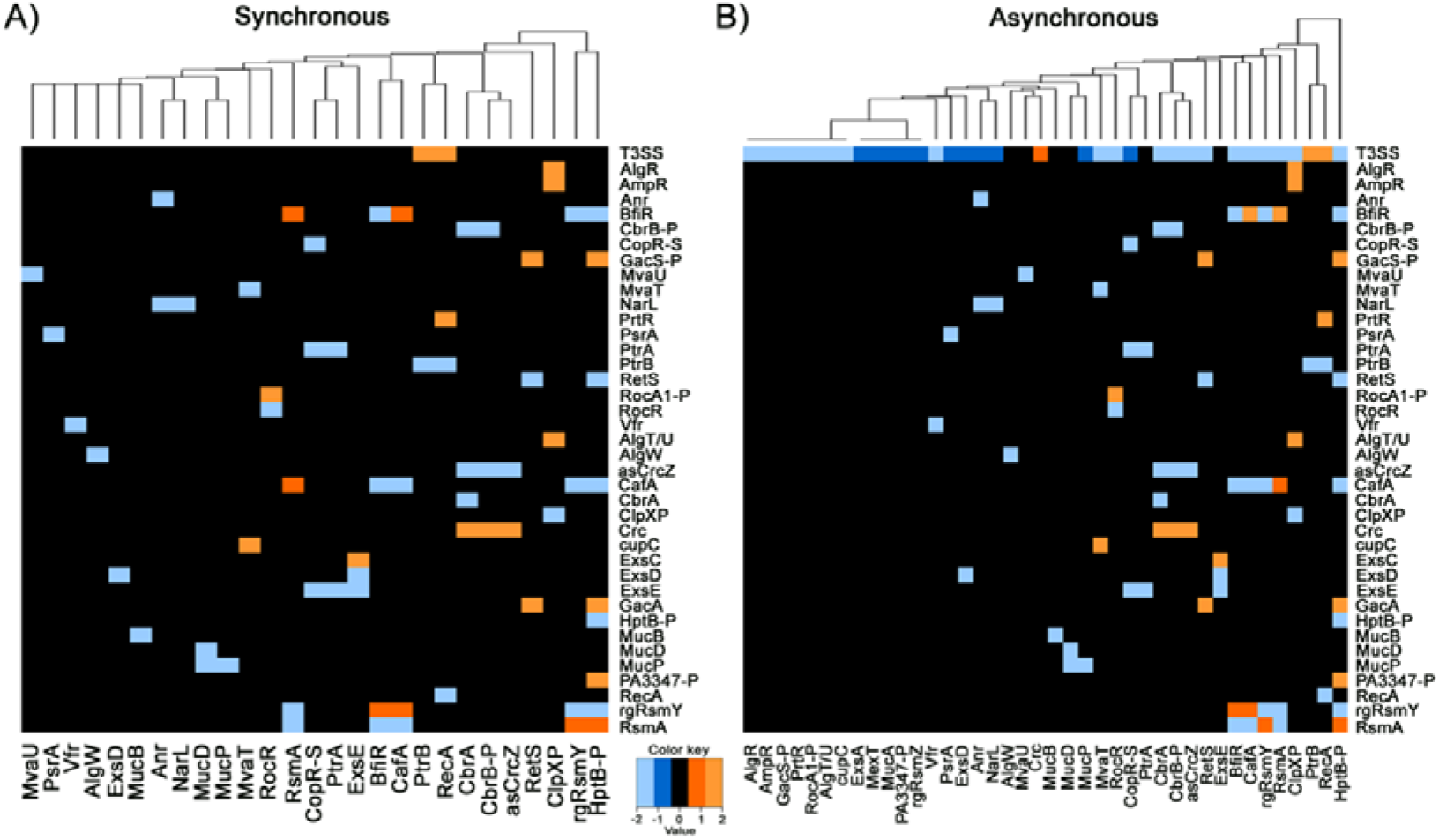
Activation patterns of the perturbed T3SS network. Effect of single node deletions from the T3SS network under the A) synchronous and B) asynchronous updating modes with all the initial conditions ON. The heat map colour indicates if the node knockout entails a lower (blue) or higher (orange) activation of a component.

### Chronic virulence networks

While the attractors of some of the acute virulence networks led to sustained activation of the virulence factor, none of the chronic networks showed sustained activation of the virulence factor when all the initial conditions were ON. Furthermore, most of the chronic networks had their virulence factor node under tight repression. Although we analysed all the chronic networks, in the following section we describe only a selection of them. See Suppl File 3 for additional information on other identified relevant networks not discussed in this main text.

- *Strictly repressed networks with moderate virulence factor activation*

To identify which nodes are key for the network functionality, we studied the network activation patterns upon node deletions and reduction of node activity. The network for the B-lactam resistance (Suppl File 1) was under strict repression, with only deletion of the initial condition ClpXP leading to higher activation of the resistance in an oscillatory pattern under both updating modes (Fig 5A). When we examined the effect of pairwise combinations of node deletions, we saw that only those involving ClpXP led to the resistance node having 50% of activation. We also found that only the 50% reduction of activation of ClpXP led to a change in the network activation pattern. Similarly, the attractors of the large biofilm formation network (Suppl File 1) were for biofilm to be OFF under both updating modes, with only the deletion or reduction of activity of the initial condition Fur leading to an increase in biofilm activation (Suppl Fig 6).

**Figure 5.**
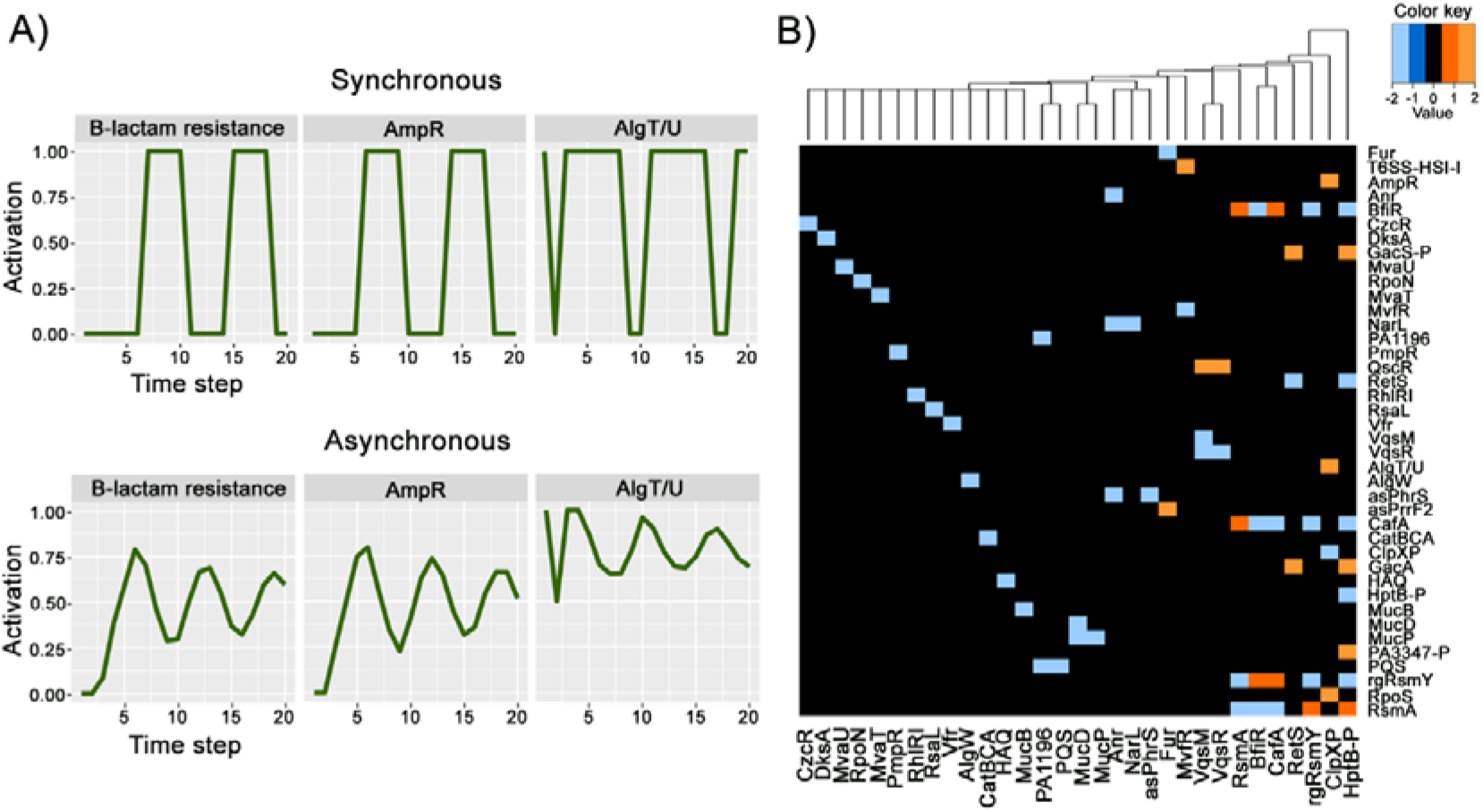
Activation patterns of chronic virulence networks. A) Activation pattern of the B-lactam resistance network with the initial condition ClpXP OFF under the synchronous (top row) and asynchronous (bottom row) updating modes. B) Effect of single node deletions on the T6SS-HSI-I network. The heat map colour indicates if the node knockout entails a lower (blue) or higher (orange) activation of a component.

The T6SS-HSI-I network (Suppl File 1) also tended towards a sustained inactivation of the virulence factor. Under both updating modes, T6SS-HIS-I was only briefly ON in *t*(1-3), and the 8 attractors had T6SS-HIS-I OFF. In our single node deletion analyses, we saw that only the deletion of MvfR led to higher activation of the factor in an oscillatory pattern (Fig 5B, Suppl Fig 7). These results suggest that a network wiring in which one key node can activate the virulence factor in an oscillatory pattern is a useful strategy exploited by *P. aeruginosa* to activate or repress its virulence rapidly, because as the network is not fixed on one state, it can easily favour one state over the other.

- *Non-strictly repressed networks with moderate virulence factor activation*

We hypothesized that the final state to which a virulence network tends to evolve is influenced by stochasticity in their regulations. To examine this, we performed all the simulations in all our models under the deterministic synchronous and the stochastic asynchronous updating modes. We found some networks in which there was a difference in the activation patterns between the updating modes. For example, we saw that the network of the fluoroquinolone resistance (Suppl File 1) under the synchronous mode had two attractors, both with the resistance node ON, and that the resistance node had sustained activation because the oscillatory phases of MexEF-Oprn and MexT were unaligned (Suppl Fig 8A). On the other hand, the asynchronous mode showed that the fluoroquinolone resistance node had 50% sustained activation (Suppl Fig 8B) and two attractors, one with the resistance node ON and one with it OFF. Furthermore, in the synchronous mode, the list of single node deletions leading to less fluoroquinolone resistance had MexS, MexT, MexEF-OprN, and ClpXP (Fig 6A), while this list in the asynchronous mode did not include MexEF-OprN (Fig 6B). To deeper examine the difference between the updating modes, we evaluated the asynchronous mode with all possible combinations of active initial conditions of the network being ON. Out of the 31 combinations, 15 led to an inactive resistance node, 7 to a ~0.3% activation, and 9 to a 50% activation (Suppl File 4). Thus, the synchronous mode hid the details of the overall tendency of the network to repress the resistance node. Taken together, these results highlight the importance of incorporating stochasticity into simulations of biological networks.

**Figure 6.**
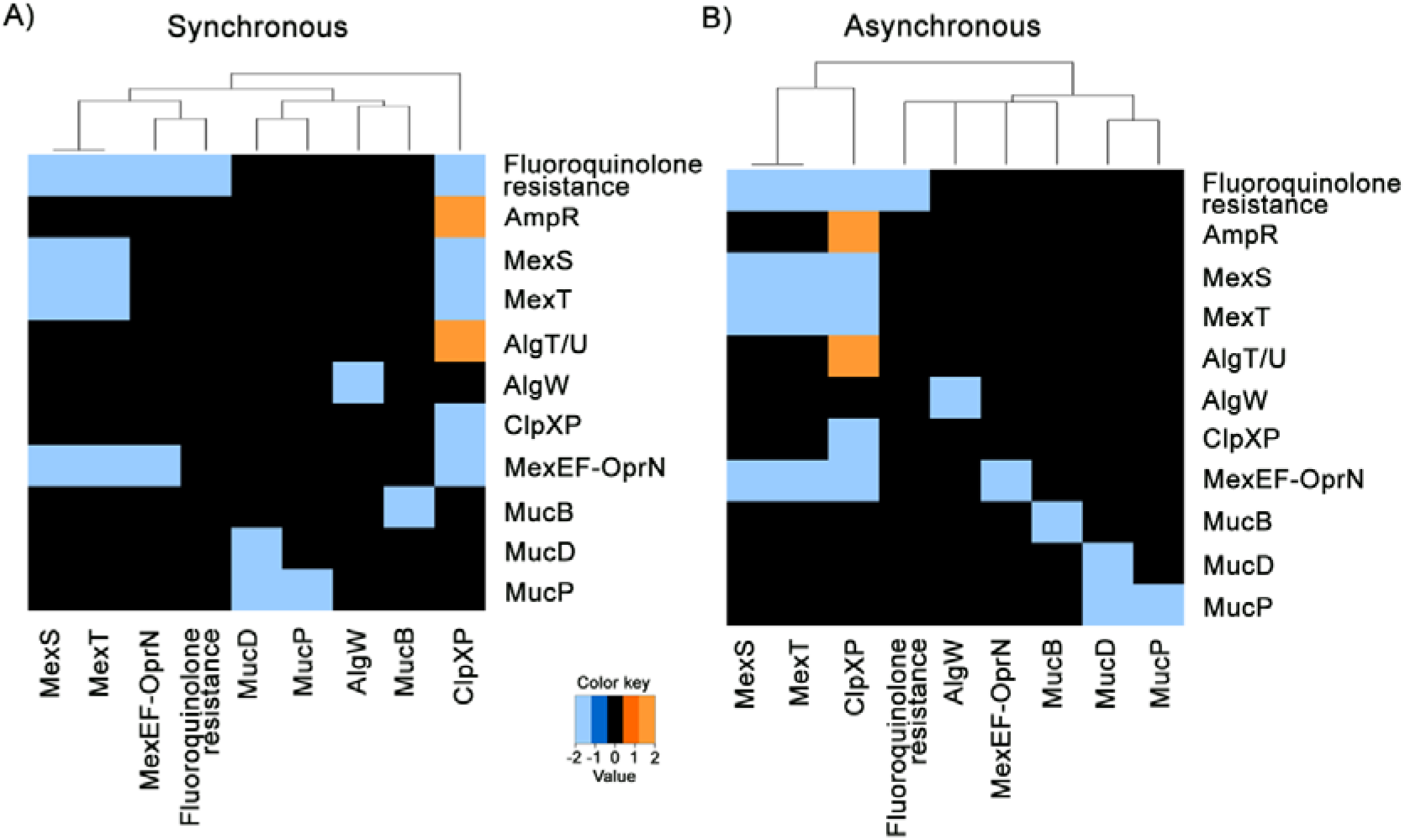
Effect of single node deletions from the fluoroquinolone resistance network. A) Under the synchronous and B) asynchronous updating modes. The heat map colour indicates if the node knockout entails a lower (blue) or higher (orange) activation of a component.

## Discussion

The QS and virulence networks function under a plethora of regulators at the transcriptional, translational, and post-translational levels. Thus, there could be several unexpected bacterial system responses that a therapy strategy targeting them could face [33]. Our models provide a computational tool to evaluate the inherent dynamicity, robustness, and stochasticity characteristics of the networks. These characteristics are important to identify which are the alternative states that *P. aeruginosa* networks could reach as a result of treatments targeting them.

### QS network suitability as drug target

QS-deficient variants carrying a mutation in *lasR* are frequently isolated from acute and chronic infections [26]. These mutants represent social cheaters, which benefit from the common goods produced by the non-mutants. However, other mutations can arise in the *lasR* mutants to turn the cheaters back into co-operators. Such a compensatory mechanism can involve the up-regulation of *rhlI* [34][27], so that it has been suggested to target RhlR for treatment [27], [35]. In our model, when the Las system repressor rsaL was ON and there was the minimum Las Al to activate LasREG only for a limited number of steps before it was overtaken by rsaL, Rhl_REG_ was able to reach sustained activation (Fig 2A). As oxidative stress activates rsaL, this result suggests that even if the immune system or drugs exert an oxidative stress to kill *P. aeruginosa,* if there is enough Las AI in the environment, *P. aeruginosa* can keep responding with virulence through the Rhl_REG_. This supports the idea of targeting with drugs the Rhl system instead of the Las system.

Compensatory mutations warn about the unaccounted side effects of targeting the QS system that could give rise to loss of treatment sensibility. Nevertheless, *P. aeruginosa* strains deficient for both *las* and *rhl* and with no compensatory spontaneous mutations have been identified [36], encouraging the development of treatments where more than one element of the QS is targeted. When we modelled the inactivation of both Las_REG_ and Rhl_REG_, we found that the PQS system could still be transiently activated if enough PQS AI was in the environment (Suppl Fig 2). As the PQS system influences the production of various virulence factors [11], [37], we investigated the effect of node deletions and AI availability on the PQS_REG_ activation. We found that the presence of PQS AI allowed the system to withstand node deletions (Fig 2C). Taken together, the model correctly represents the ability of *P. aeruginosa* to modulate the expression of PQS_REG_, because PQS is an AI that if present in too large amounts is detrimental to the pathogen, while when present in moderate amounts, it is used for stress response [38].

A previous study on the Boolean models of three QS systems (Las, PQS and Rhl) examined at which step the Las_REG_, Rhl_REG_, and PQS_REG_ nodes turn ON. The study concluded that the regulation of timing played a secondary role in the QS system operation, with the wiring between the nodes playing the dominant role, so that developing drugs targeting these interconnections is a good strategy. In our study, we re-evaluated this hypothesis in light of the finding of the new IQS system and the availability of new more powerful software for BM [20]. We found that modifying the times of the modulations between the Las and Rhl systems led to discrepancies between the results of the updating modes. This highlights that the timing of the activity of the effector nodes, and not only the node connections, has a relevant impact on the system (Fig 2D, Suppl Table 1). Notably, for all the perturbations, both updating modes led to the same patterns. In principle, this would suggest that the wiring of the network itself is what gives the resilience to the QS in *P. aeruginosa.* However, these patterns are from the final states of the system. When examining the systems through time, we discovered properties unique to the asynchronous mode, such as longer activation of a system or a range of percentages of activation.

We conclude that the utility of QS as a druggable target is relevant, although the network inherent dynamicity needs to be carefully considered [5]. More importantly, its interconnectivity with other virulence factors should also be evaluated. As observed, deletion of the MvfR node, part of the PQS system, leads to activation of the T6SS-HIS-I [39] (Fig 5B), which is in turn required for biofilm formation [40].

### Network inherent dynamicity and robustness

The attractors of a network represent the long-term behaviour of the system, thus analysing them provides insights into the networks preference of states. For example, if a network has only one attractor where the relevant node is OFF (e.g. a virulence factor), it suggests that the network will be less prone to turn the node ON. Furthermore, the perturbation analyses and the examination of the effect of the time lag for the effective activity of a regulator node allow for the identification of regulatory interactions that are key to the dynamicity and robustness of the system.

In terms of dynamicity due to the network’s attractor space, we uncovered that an oscillatory behaviour is a useful strategy to activate or repress the production of a virulence factor whenever required, likely as a strategy for a rapid stress response. For example, we found this oscillatory behaviour in the elastase network (Suppl Fig 5). Importantly, experimental validation of this *in silico* observation has been previously reported, showing that when *P. aeruginosa* is in a metal ion-deficient environment there is a low level of produced elastase, but as soon as metal ions are supplemented, high levels of elastase are detected [41].

Opposite to networks that harbour a range of available attractors that make their responses more dynamic, other networks are under very strict regulation, so that they only tend to one state towards which only few paths lead to. An example of such a robust network is that of flagella, in which BifA is the key node able to modify the network’s attractor (Suppl Fig 4C), as previously experimentally demonstrated [42]. It is expected that flagella is under such tight control, as transitions from motility to stationary lifestyle and vice versa have a strong impact in the *P. aeruginosa* metabolism. This lifestyle largely depends on whether the pathogen is in an acute infectious mode, where motility is needed for invasion, or a chronic infectious mode, where a stationary behaviour is needed for biofilm formation [43].

The model for biofilm formation showed a very robust repressive behaviour (Suppl Fig 6). However, it should be considered that the large and complex biofilm biological system includes several environmental conditions influencing the different stages of biofilm formation, and more elements which are not included in our model can be implicated [44], [45]. Our reconstructed network captures the overall system structure without dividing it by the biofilm formation stages, which likely need to be modelled separately in order to provide a clearer picture. Nonetheless, the model correctly illustrates that the system heavily controls biofilm formation.

### Network inherent stochasticity

Mathematical modelling of biological systems provides valuable insights into the complexity of biological systems, as they dynamically evolve through time and may do so non-linearly. In our study, the stochastic (asynchronous) mode provided deeper insights into the behaviour of some of the networks than the deterministic (synchronous) mode. This observation is supported by other studies that demonstrate that stochasticity plays a role in processes such as bacterial gene expression [46] and chemotaxis [47].

Notably, the asynchronous mode on the pyoverdine production network showed that a very small percentage of the simulations did not produce pyoverdine. This suggest that, due to stochasticity, members of a *P. aeruginosa* population could arise as “cheaters” that benefit from the pyoverdine produced by the majority of the population, while relieving themselves from the metabolic burdens of its production. These cheaters could then have a fitness advantage if the conditions changed, preventing the infection to be completely eradicated. Such a pyoverdine production cheating behaviour has been experimentally studied in detail in *P. aeruginosa,* identifying that there is both selection for wild-type production and for cheating, with the mean fitness of the population determined by the difference of the benefits and the cost of pyoverdine production [48], [49].

We also observed a stochasticity effect in the T3SS network, where although most of the attractors of the system had T3SS OFF, few cases arose where it was ON (Suppl Table 6, Fig 4). Although the T3SS formation is a complex process [50], this result suggests that at least a small portion of the *P. aeruginosa* population is able to utilize this system to initiate infection. It could be that there are regulatory elements or other environmental signals not accounted for in the model reconstruction that lead to such behaviour. For example, within populations of *Yersinia pseudotuberculosis* where the members have the same or closely similar genotypes, T3SS expression is associated with proximity to host cells [51]. Overall, our modelling results highlight that further research is needed to elucidate the complete molecular network leading to the regulation of virulence factors.

It is interesting that even though this stochasticity component could derive from regulatory elements and signals not accounted for in the network reconstructions, the models are nonetheless able to reproduce its effect. For example, fluoroquinolone resistance in *P. aeruginosa* has been shown to be influenced by the differential expression of hundreds of genes involved in other disparate processes, such as bacteriophage-like pyocins and regulons controlled by the LexA-like SOS response [52], [53]. Notably, a stochasticity effect was observed in our model of fluoroquinolone resistance analysed under the asynchronous mode (Fig 6).

### Repressive nature of chronic virulence networks

Interestingly, while the acute virulence networks present a range of patterns that can lead to either activation or repression of the virulence factor, the patterns of the chronic networks can only be classified by their repression strategy. Our model of beta-lactam resistance correctly represents the experimental observation that the expression level of beta-lactamase is usually low [54] (Fig 5A). Notably, the molecular basis for the induced production of beta-lactamase, the enzyme that enables beta-lactam resistance, in Gram-negative bacteria is not fully clear [55], thus our results contribute to the current efforts in understanding their regulatory mechanisms [56].

## Conclusions

The use of computational models within the microbiology research field has proven useful for the understanding of bacterial pathogenicity. In this study, we use BM to address the challenge of integrating different metabolic signals and regulators into a modelling framework that sheds light into new details of the adaptability mechanisms of *P. aeruginosa,* providing a deeper understanding of its pathogenicity. Importantly, the models we developed can serve as tools for future studies to integrate newly identified elements connected to the network [57], or to eliminate false-positive interactions from the literature-based network and predict new interactions [58].

Importantly, our results suggest that prior to any drug development efforts, the effects of targeting the chosen element (e.g. a node from the Rhl system) should be carefully evaluated as the targeted system can have unexpected behaviours that lead it back to its attractor state where the virulence factor is active. Notably, these insights obtained through our BM had not been previously unveiled by other metabolic modelling techniques, highlighting the importance of using various modelling techniques to support and guide drug design and experimental work [59].

## Acknowledgements

We thank Dr. Vanessa Phelan, Dept. of Pharmaceutical Sciences, University of Colorado, for her insights into the metabolism of *P. aeruginosa.* We are also thankful to the Birmingham Environment for Academic Research (BEAR), particularly Radoslaw Poplawski, for providing the needed computational resources and IT support in the development of this project. This research has been funded by the European Union’s Horizon 2020 Research and Innovation Programme under the Marie Sklodowska-Curie Grant Agreement No 836384.

